# Closed-loop Neuroscience of brain rhythms: optimizing real-time quantification of narrow-band signals to expedite feedback delivery

**DOI:** 10.1101/2019.12.18.880450

**Authors:** Nikolai Smetanin, Anastasia Belinskaya, Mikhail Lebedev, Alexei Ossadtchi

**Affiliations:** Center for Bioelectric Interfaces, Higher School of Economics, Moscow, Russia, 101000

**Keywords:** Closed-loop neuroscience, neurofeedback, brain rhythm, envelope, instantaneous phase, feedback latency, optimal filtering, Hilbert transform

## Abstract

Closed-loop Neuroscience is based on the experimental approach where the ongoing brain activity is recorded, processed, and passed back to the brain as sensory feedback or direct stimulation of neural circuits. The artificial closed loops constructed with this approach expand the traditional stimulus-response experimentation. As such, closed-loop Neuroscience provides insights on the function of loops existing in the brain and the ways the flow of neural information could be modified to treat neurological conditions.

Neural oscillations, or brain rhythms, are a class of neural activities that have been extensively studied and also utilized in brain rhythm-contingent (BRC) paradigms that incorporate closed loops. In these implementations, instantaneous power and phase of neural oscillations form the signal that is fed back to the brain.

Here we addressed the problem of feedback delay in BRC paradigms. In many BRC systems, it is critical to keep the delay short. Long delays could render the intended modification of neural activity impossible because the stimulus is delivered after the targeted neural pattern has already completed. Yet, the processing time needed to extract oscillatory components from the broad-band neural signals can significantly exceed the period of oscillations, which puts a demand for algorithms that could minimize the delay.

We used EEG data collected in human subjects to systematically investigate the performance of a range of signal processing methods in the context of minimizing delay in BRC systems. We proposed a family of techniques based on the least-squares filter design – a transparent and simple approach, as it required a single parameter to adjust the accuracy versus latency trade-off. Our algorithm performed on par or better than the state-of the art techniques currently used for the estimation of rhythm envelope and phase in closed-loop EEG paradigms.

## 1 Introduction

Investigations of neural oscillations have a long history and continue to be an area of intensive research, particularly when such neuroimaging techniques are used as noninvasive electroencephalography (EEG) and magnetoencephalography (MEG), and invasive electrocorticography (ECoG) and stereo EEG (sEEG).

A plethora of experimental paradigms and relevant analysis methods have been developed for dealing with specific types of neuronal oscillations, including the methods for their induction and suppression [1, 2]. These paradigms fall in one of two categories. In the first category of studies, changes in neural oscillations are investigated that are induced by a variety of stimuli; the stimuli are presented without a consideration of the ongoing brain activity. In the second category [3], a closed-loop design is implemented where the stimuli are selected based on the characteristics of the ongoing brain activity.

Below, we describe the studies from the second category where the closed loop is formed from neural oscillatory activity. We refer to this experimental approach as brain rhythm contingent (BRC) paradigm.

### 1.1 Brain rhythm contingent paradigms

As shown in Figure 1A, the BRC paradigm operates through three steps: data acquisition, data processing, and stimulus generation. During the data acquisition step, brain activity is measured, usually with multiple spatially distributed electromagnetic sensors, and streamed to a computer. During the data processing step, a computer routine handles the multichannel data in real-time to extract the parameters of oscillatory activity, typically amplitude and phase. Lastly, during the stimulus generation step, these parameters are converted into a feedback delivered to the brain either directly, using stimulation applied to the nervous tissue (also called neuromodulation), or through natural senses: vision, hearing or touch. The feedback could act in a subtle way by modulating the parameters of an already ongoing stimulation (direct or through natural senses) or by contributing to the algorithm that selects a stimulus from a set of discrete possibilities.

**Figure 1:**
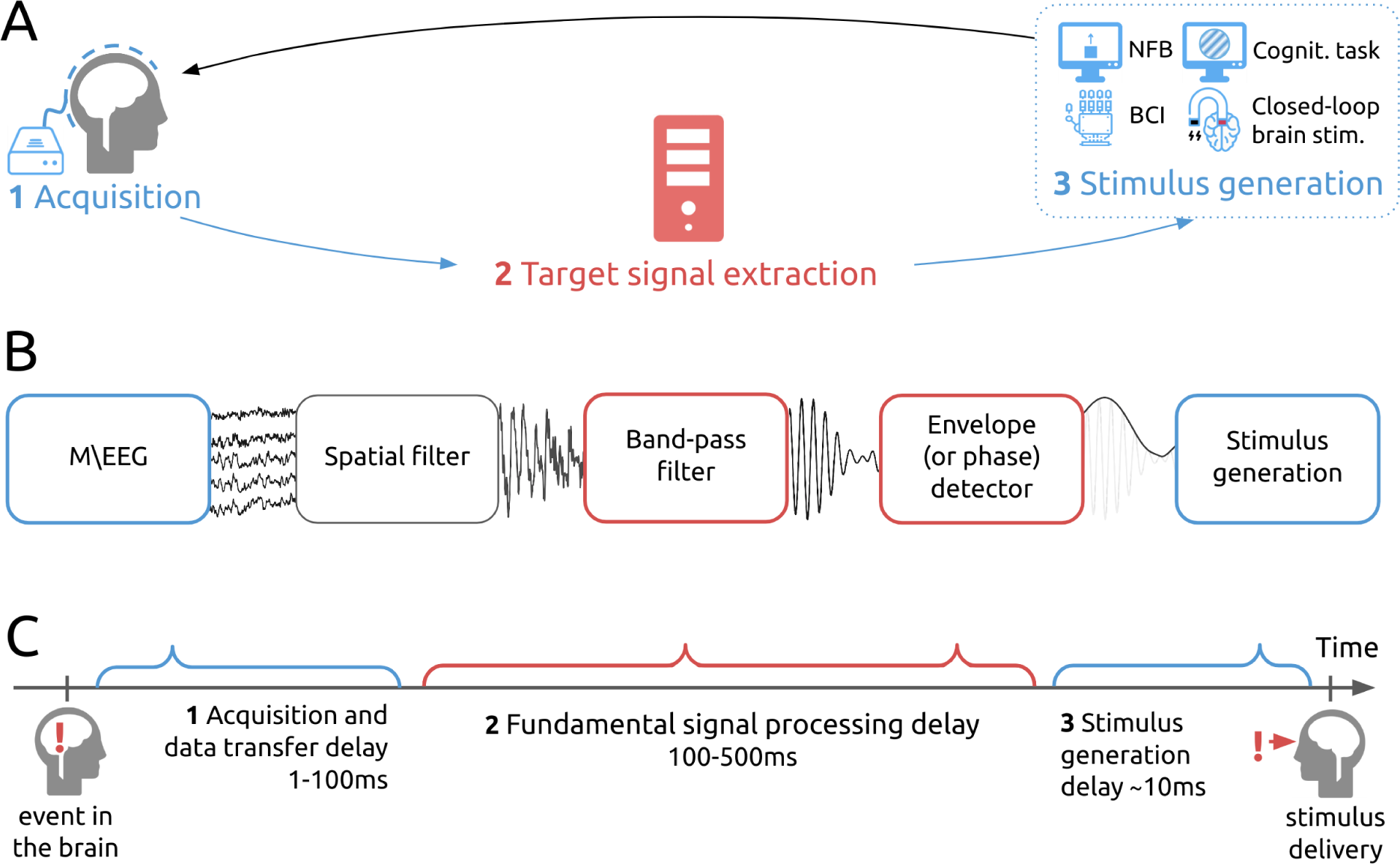
Schematics of the BRC paradigm. A. A diagram depicting signal flow in a closed-loop system. B. Signal processing pipeline. C. The sources of delays mounting to the total latency of the BRC system. Technical and fundamental sources of the delay are marked in blue and red, respectively.

Many implementations of BRC paradigm have been developed, which allow implementation of a variety of goal-directed behaviors dependent on a closed-loop design [3]. The most distinct paradigms are: neurofeedback (NFB), brain computer interface (BCI), closed-loop brain stimulation, and brain state-contingent stimulus delivery.

The NFB is a form of biofeedback that enables subjects with the capacity to monitor and control their own brain activity [4]. With NFB, subjects gain access to the neural signals of different brain structures, and learn to modulate them in specific ways [5], [6], [4], [7]. NFB approach is used as a therapy for neurological disorders [8], [9] and as cognitive enhancement therapy [10]. Steps of NFB operation include extraction of neural features of interest, their transformation using case-specific algorithms, and generation of sensory feedback delivered to the subject.

*BCIs* operate very much like NFB, with neural activity being recorded, processed, and directed to an external device that provides some sort of feedback to the subject. Yet, the emphasis here is not on the feedback per se but on the brain control of the device, which serves some useful purpose, for example a computer cursor or a prosthetic limb [11]. Clinically relevant BCIs are intended for functional restoration and rehabilitation of patients with neurological disabilities. For example, an EEG-based, motor-imagery BCI that controls a hand exoskeleton aids in rehabilitation of stroke patients [12], [13] by facilitating Hebbian plasticity that occurs owing to synchronization of cortical modulations with proprioceptive feedback caused by exoskeleton movements [14].

In BRC paradigms with *closed-loop brain stimulation*, parameters of neural oscillations, most often instantaneous amplitude and phase, affect the characteristics of stimulation directly applied to the brain [15]. Electromagnetic devices are commonly used to deliver the stimulation, which can be invasive, such as deep brain stimulation (DBS) [16], or noninvasive, such as transcranial brain stimulation (TBS) [17]. Thus, EEG oscillatory patterns have been used to control such types of TBS as transcranial magnetic stimulation (TMS) [18] and transcranial alternating current stimulation (tACS)[19]. BRC paradigms with stimulation through normal senses can be considered a type of closed-loop brain stimulation, as well, for example, stimulation with continuously flashing visual stimuli[20].

BRC paradigms have gained popularity, where cognitive processes are investigated under experimental conditions with *brain rhythm-contingent stimulus delivery*. This is because the brain handles cognitive tasks differently depending on the brain oscillatory patterns taking place just before the task onset [21, 22, 23, 24, 25, 26, 27]. Accordingly, it is of interest to create an experimental paradigm, where the task starts only after a particular oscillatory pattern is detected. Such brain rhythm-contingent stimulus delivery can drastically reduce experimental time and minimize participants’ fatigue, particularly when the neural patterns of interest are represented by short-lived bursts of activity [28] or desired brain states[29].

### 1.2 Signal processing and latency in BRC paradigms

Despite the conceptual differences between the four applications of the BRC paradigm described above, their signal processing pipelines are quite similar and can be summarized as shown in Figure 1B.

In this paper, we consider the BRC paradigms based on EEG and MEG recordings. The major advantage of these imaging techniques over those that employ metabolic (e.g., positron-emission tomography) or blood oxygenation level-dependent (e.g., functional magnetic resonance imaging) measurements is their unsurpassed millisecond-scale temporal resolution. This fine temporal resolution allows exploring fine rhythmic structure of brain activity and, in principle, enables nearly instantaneous interaction with the brain circuits. Yet, as explained in detail below, time-lags of both technical and fundamental nature occur during the online extraction of oscillatory parameters from the ongoing brain activity. These lags vary from one implementation to another and can be significantly reduced with the use of appropriate signal processing methods.

For a BRC paradigm to be efficient, it needs to be temporally specific, that is feedback should be issued when it can affect the targeted neuronal activity. Temporal specificity of a BRC system is characterized by an overall delay between the onset of the neural event of interest and the time when the participant receives the stimulus corresponding to this event (Figure 1C. This delay, called the overall BRC system latency, incorporates time-lags related to different factors, some of them technical (i.e, software and hardware delays) and some fundamental (i.e. required to collect a sufficient amount of neural data).

The technical time-lags typically do not exceed 100 ms; this time is needed for hardware communication and low-level software processing. This delay can be reduced by specific hardware and software solutions. Based on our experience, it is feasible to reduce the technical delay to 20-30 ms.

The fundamental lag cannot reduced that easily because it is composed of the time needed to collect a snapshot of neural oscillatory data that could be then quantified with an appropriate algorithm like band-pass filtering followed by the extraction of information about the instantaneous power or phase of the narrow-band process. If suboptimal approaches are used for extracting power and/or phase of rhythmic neural components, the fundamental delay amounts to approximately 0.5 s, causing undesirable effects in BRC implementations.

The adverse effect of feedback delays has been reported in many NFB and the BCI studies, where participants learned to properly modulate their own brain activity, with or without a strategy recommended by the experimenter [30]. A similar problem with long feedback delays has been known from the studies on evidence-based learning. Thus, back in 1948, Grice showed that learning to discriminate complex visual patterns drastically depended on the feedback signal latency [31]. Impaired performance with delayed has been also demonstrated in the studies on motor learning e.g for the prism adaptation task [32]. According to Rahmandad et al. [27], behavioral learning is impaired when feedback delay is unknown. Moreover, elongated feedback delay decreases the sense of agency during BCI control [33], the finding also corroborated by a simulation study [34] showing that feedback delay and temporal blur adversely influence the automatic (strategy free) learning.

Temporal specificity is also an an important consideration for the experimental settings with closed-loop brain stimulation and brain state triggered stimulus delivery. Indeed, the key requirement for these methods are the accurate estimation of instantaneous oscillatory features and the timely delivery of the stimuli to efficiently interfere with the oscillatory neural patterns.

### 1.3 Low-latency method for envelope and phase detection

In the present study, we explore several approaches aimed at reducing the delay between neuronal events and the corresponding feedback in the BRC paradigm. We propose a simple method to directly control the delay. Our algorithm is based on the adaptive optimization of the complex-valued finite impulse response (FIR) filter weights to yield the desired reduction of delay. The proposed approach is capable of extracting the instantaneous power and phase of neural oscillations with the shorter latency and higher accuracy as compared to other relevant techniques. With this approach, the user can explicitly specify the desired delay and assess the corresponding accuracy of envelope estimation. These features of the algorithm allow for flexible control of the latency-accuracy trade-off in BRC applications. Strikingly, our approach can work even with negative delays, paving way for achieving predictive feedback.

## 2 Methods

Our basic assumption is that the measured neural activity *x*[*n*] is a sum of the narrow-band signal *s*[*n*] (targeted neural activity of a BRC paradigm) and background colored broad-band noise *η*[*n*].

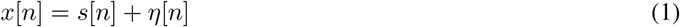

The targeted neural activity *s*[*n*] can be represented as the real part of analytic signal *y*[*n*] [35]:

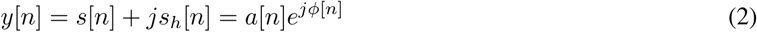

where *s*_*h*_[*n*] is the imaginary part of the analytic signal, often called “second quadrature” of the original signal *s*[*n*], *a*[*n*] is the envelope, reflecting the instantaneous power of the narrow band process at time stamp *n*, and *ϕ*[*n*] is its instantaneous phase at the time stamp *n*. Importantly, once the estimate of the complex valued analytic signal *y*[*n*] is available that corresponds to signal *s*[*n*], the envelope *a*[*n*] and the phase *ϕ*[*n*] of *s*[*n*] can be computed as

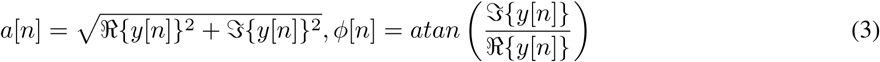

Figure 2A shows that, with these expressions, extraction of the ground-truth signal is not restricted to causal operations. This is because the discrete Fourier transform (DFT) and narrow-band Hilbert transform in the frequency domain can be applied to the entire batch of data, followed by the conversion of the real and imaginary components of the analytic signal into the ground-truth values of the instantaneous envelope *a*[*n*] and phase *ϕ*[*n*].

**Figure 2:**
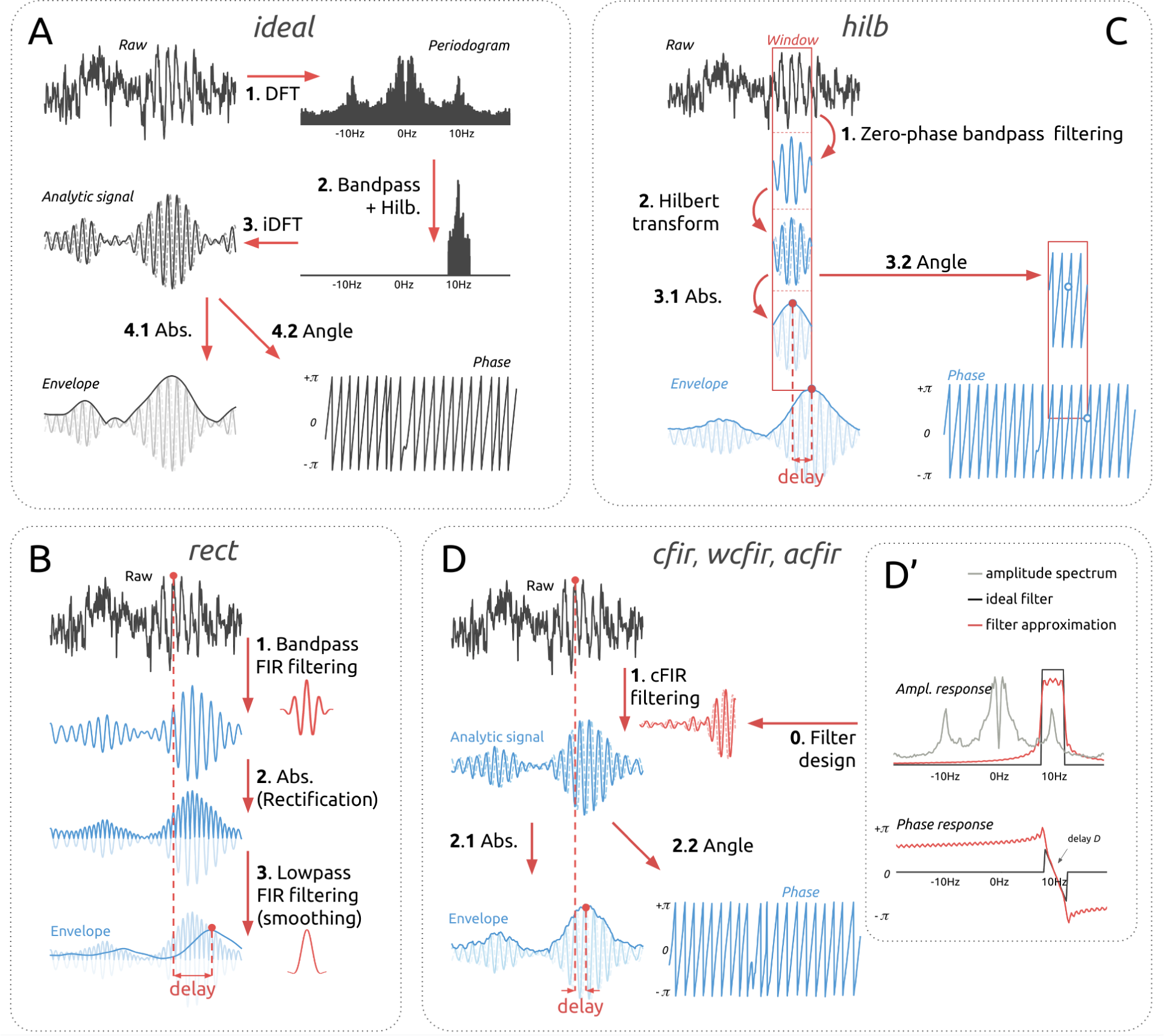
Methods for narrow-band signal envelope estimation. A. Ideal non-causal system for ground truth signal extraction. B. Envelope detector based on rectification of the band-filtered signal. C. Sliding window narrow-band Hilbert transform-based method. D. cFIR family filters that perform finite impulse response causal filter approximation of ideal non-causal systems. D’. Filter design for cFIR family filters.

In addition to the non-causal approach illustrated in Figure 2, the other panels of Figure 2 show a graphical summary of the available causal techniques that can be applied in real time to compute the envelope and instantaneous phase of a narrow-band component extracted from a broad-band signal.

### 2.1 Existing methods

For didactic purposes, we start with the most basic approaches.

#### Rectification and smoothing of the band-filtered signal (*rect*)

is the conceptually most straightforward way to estimate the envelope *a*[*n*]. In this method, low-pass filtering is performed applied to the rectified narrow-band filtered signal. This approach can be mathematically expressed as:

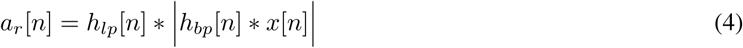

where is the convolution operator, *h*_*bp*_ is the impulse response of the band-pass filter, |⋅| denotes the absolute value (i.e., the rectification step), and *h*_*lp*_ is the impulse response of the low-pass filter that performs smoothing of the rectified signal. Without loss of generality, we can assume that both *h*_*bp*_ and *h*_*lp*_ are linear phase FIRs designed by a Hanning window method with the number of taps *N*_*bp*_ and *N*_*lp*_, correspondingly. The cutoff frequency for the low-pass filter *f*_*c*_ is set to correspond to one-half the expected bandwidth of the narrow-band process, i.e. *f*_*c*_ = (*f*_2_ − *f*_1_)/2. FIR filters have a linear phase and therefore the total delay *D* and the number of taps in the individual filters *N*_*bp*_ and *N*_*lp*_ have the following interrelationship:

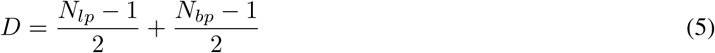

In order to ensure that the maximum performance is achieved for a given group delay value *D*, we used grid search over variables *N*_*bp*_ and *D* in our comparative analysis. The parameter *N*_*lp*_ was determined from the formula 5. Since *N*_*bp*_ and *N*_*lp*_ are positive, this method can estimate the envelope values only with a positive delay, which corresponds to the count of signal samples taken from the past.

#### Sliding window narrow-band Hilbert transform (*hilb*)

is the second most commonly used method that is based on the use of the analytic signal *y*[*n*] computed using Hilbert transform [35]. There are various implementations of this approach. In the current work we resorted to the use of the windowed DFT. DFT is calculated on each window of length *N*_*t*_ which is zero-padded to the length of *N*_*f*_ samples. Next, the coefficients corresponding to the negative frequency values and those in the positive frequency semi-axis that fall outside the band of interest are zeroed out. The DFT coefficients within the band of interest are doubled. Then, the inverse DFT is performed and *N*_*t*_ − *D*-th element of the resultant complex valued sequence is used as an estimate of the analytic signal with delay *D*. This way, two operations are performed simultaneously: band-pass filtering and extracting the analytic signal that is then used to estimate the envelope. This algorithm is illustrated in Figure 2.

In matrix representation and using temporal embedding to form vector, **x**[*n*] this can be written as:

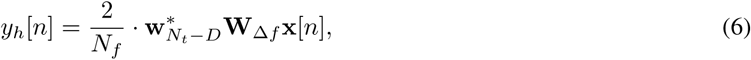

where vector **x**[*n*] contains the last *N*_*t*_ samples of *x*[*n*], i.e. **x**[*n*] = [*x*[*n*], *x*[*n* − 1],… *x*[*n* − *N*_*t*_ + 1], **W**_∆*f*_ is the *N*_*f*_ -by-*N*_*t*_ modified DFT matrix with zeros on the *k*-th row for *k* outside of [*f*_1_*N*_*f*_/*f*_*s*_, *f*_2_*N*_*f*_/*f*_*s*_] range corresponding to the physical band of interest ∆*f* = [*f*_1_, *f*_2_] and 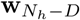 is the (*N*_*h*_ − *D*)-th row of the DFT matrix.

The parameters to be optimized for this method are window length *N*_*t*_ and zero-padded length of the signal *N*_*f*_ which is used to perform the DFT. The overall delay of this method is explicitly determined by parameter *D*.

#### Sliding window Hilbert transform with AR prediction of the narrow-band filtered signal (*ffiltar*)

was proposed by Chen et al. [36] and applied practically [18]. In this method, the sliding window vector **x**[*n*] containing *N*_*a*_ last samples is forward-backward band-pass filtered. Then, flanker *N*_*e*_ samples are truncated to eliminate edge artifacts, and 2*N*_*b*_ samples are forward predicted by using an AR model fitted to *N*_*a*_ − *N*_*e*_ samples. Finally, Hilbert transformation of prediction is used to estimate the analytic signal value in the middle of the predicted range. In the original work, this value was used to determine the current phase and time for stimulation. Here we used this technique as a benchmark for our and other methods being compared but only at the processing latency *D* = 0, that is the latency this approach was originally designed for.

Using matrix notation, we can formalize this method as follows.

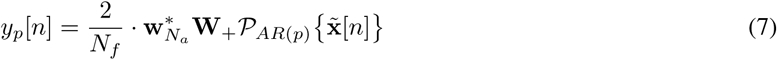

where 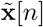 contains forward-backward filtered last *N*_*a*_ samples of the *x*[*n*], 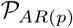 denotes AR model based prediction operation and adds 2*N*_*e*_ predicted samples by using a *p*-th order AR model, **W**_+_ is the *N*_*f*_-by-(*N*_*a*_ + *N*_*e*_) modified DFT matrix with zeros on the rows corresponding to the negative frequencies and 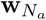 is the (*N*_*a*_)-th row of the DFT matrix. As the narrow-band filter for the forward-backward filtering part, we use Butterworth filter of the order *k* as suggested in [36].

One of the disadvantages of this approach is that it has multiple parameters that need to be tuned to achieve the optimal performance. To attain the best performance in this study, we searched over the parameter grid composed of the following variables: AR order *p*, number of edge samples *N*_*e*_, and Butterworth filter of the order *k*.

### 2.2 Proposed method

In order to build the analytic signal *y*[*n*] that corresponds to the narrow-band signal *s*[*n*] extracted from the noisy broad-band measurements *x*[*n*], one can apply the ideal narrow-band Hilbert transform filter [37, 38]. The complex-valued frequency response of this combined filter can be defined as:

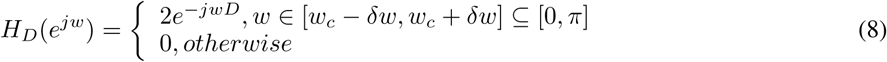

where *δw* = 2*πδf* is half of the pass band width, and *D* is the group delay measured in samples. Strictly speaking, for any finite delay *D* this filter is non-causal and cannot be applied in real time. To reconstruct the analytic signal causally, one can find a causal complex-valued finite impulse response (cFIR) filter **b** = {*b*[*n*]} of length *N*_*t*_ that approximates the ideal complex valued frequency response *H*_*D*_(*e*^*jw*^) [39]. This filter can be then applied in real-time as *y*_*c*_[*n*] = *x*[*n*] *∗ b*[*n*], the procedure that incurs a fixed processing delay of *D* samples.

Causal complex valued FIR *b*[*n*] can be found by solving the least squares optimization problem. Various definitions of the cost functions lead to different filters.

#### Frequency domain least squares (*cFIR*)

is the first and most straightforward approach (denoted F-cFIR). The least squares filter design strategy consists in finding the complex valued vector of the cFIR filter weights **b** of length *N*_*t*_ that minimizes the *L*_2_ norm between the cFIR filter frequency response obtained by the DFT and the discrete appropriately sampled version **h**_*D*_ of an ideal response *H*_*D*_ in the frequency domain. To increase the frequency resolution, we use truncated *N*_*f*_-samples DFT-matrix **W** with dimension *N*_*f*_ × *N*_*t*_(*N*_*f*_ ≥ *N*_*t*_). The use of transform matrix-based formulation of the DFT in this case is equivalent to the DFT of *N*_*t*_ samples long vector zero-padded to length *N*_*f*_. Taking in account the fact that due to orthogonality **W**^**H**^**W** = *N*_*f*_**I** the solution **b**_*F*_ of the normal equation for the optimization problem (2.2) can be found by a simple inverse DFT (9) of the desired complex valued characteristics of the narrow-band Hilbert transformer.

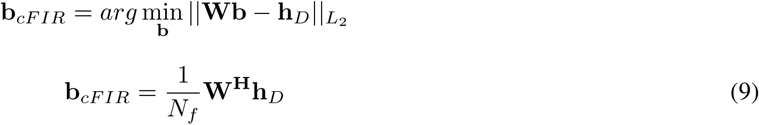

The last formula 9 is equivalent to expression 6 but can be used with negative delays, *D* ≤ 0. This simple method, however, does not take into account the second order frequency domain statistics of the target signal and could be further improved. Note that the cFIR approach with parameters *N*_*t*_, *N*_*f*_ and *D* ≥ 0 matches the sliding window narrow-band Hilbert transform approach with the same parameters *N*_*t*_, *N*_*f*_ and *D*.

#### Frequency domain weighted least squares (*wcFIR*)

is the method that follows optimal filter design ideas, where power spectral density of the input signal *x*[*n*] are used as weights. We thus formulate the weighted frequency domain least squares design technique (denoted wcFIR) via optimization problem (2.2) whose solution can be found by solving the normal equations (10):

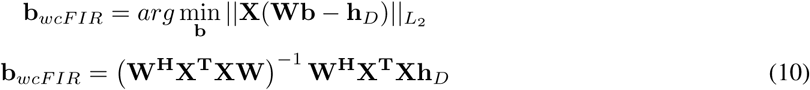

where **X** is the diagonal matrix formed from the square roots of the power spectral density magnitudes of *x*[*n*]. The temporal dimension of **W** is set according to the specified delay *D* by cropping **W** from the full-blown twiddle-factor matrix. Therefore only **W**^**H**^**W** = *N*_*f*_**I** holds true while **WW**^**H**^ ≠ *N*_*f*_**I**. This way, at the optimum ||**Wb** − **h**_*D*_|| ≠ 0 and therefore **b** is just an approximation of the ideal filter and can be computed even for negative *D*.

Panels D and D’ of Figure 2 illustrate these two approaches. The delay *D* corresponds to the slope of the phase response within the pass-band and theoretically can be set to an arbitrary value. Then, the optimization procedure aims at finding such complex vector of cFIR filter coefficients **b** that both ideal phase response and an ideal amplitude response are approximated sufficiently and accurately. We do so in the frequency domain by analytically solving the least squares or the weighted least squares problems.

Conceptually, having in mind the two tasks of optimal envelope and instantaneous phase estimation we could have formulated the two separate optimization problems and used two different sets of weights implementing two different band-pass complex-valued filters delivering optimal accuracy in estimation of envelope and phase approximation with the specified delay. In this case, however, due to non-linearity of the target functional, we would have to perform an iterative optimization in order to find the optimal FIR filter weights vector **b**.

#### Time domain least squares

is the last approach from this family (denoted tcFIR) that is based on minimization of the squared distance in the time domain between the complex delayed ground truth signal *y*[*n − D*] and the filtered signal *x*[*n*] *∗ b*[*n*]:

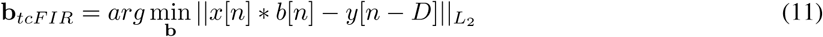

In this case, the ground truth signal *y*[*n − D*] is obtained non-causally from the training samples via an ideal zero-phase Hilbert transformer (8). According to Parseval’s theorem, this approach is equivalent to the wcFIR approach. However in contrast to the frequency domain formulation, it allows for implementation of recursive schemes for solving (11) and therefore may potentially account for non-stationarity in the data. One of the most straightforward approaches is to use recursive least squares (RLS) [40], to update filter coefficients on the fly.

### 2.3 Methods comparison

#### Data and preprocessing

We compared the described methods using resting state EEG data recorded from 10 subjects during NFB training sessions. EEG recordings were performed using 32 AgCl electrodes placed according to a 10-20-system with the ground electrode at AFz position and reference electrodes on both ears. The impedance for all electrodes was kept below 10 KOhm. The signal was sampled at 500 Hz using the NVX-136 amplifier (Medical Computer Systems Ltd.) and bandpass-filtered in 0.5-70 Hz band. These *preprocessing* filters incurred overall delay of no more than 10 milliseconds in the bandwidth of interest (8-12Hz).

For each subject, we used 2 minutes of resting state recordings. The first minute of the data – for training or the parametric grid search and the second minute is used for testing the performance. To eliminate eye artifacts we performed independent component analysis (ICA) on the training data, identified eye-movement related components by means of the mutual information spectrum [41] and removed from the data several components exhibiting the highest mutual information with either of the two frontal channels Fp1 and Fp2.

For the following analysis only parietal P4 channel of the cleaned data is used as feedback signal *x*[*n*] in (1).

#### Individual alpha band

We determine individual alpha range by the following procedure: estimate power spectrum using Welch method with 2 second 50%-overlap boxcar window, find the frequency *f*_0_ with maximal SNR in the 8-12Hz range, define individual band as [*f*_1_, *f*_2_] interval, where *f*_1_ = *f*_0_ − 2 Hz and *f*_2_ = *f*_0_ + 2 Hz.

#### Ground truth signal

As the ground truth signal *s*[*n*] in (1), we used non-causally computed analytic signal obtained by zeroing out the DFT coefficients corresponding to the frequencies of the individual alpha band [*f*_1_, *f*_2_] followed by the Hilbert transform. Once we have the analytic signal, we can estimate both envelope and instantaneous phase without any additional delay.

#### Performance metrics

To measure the accuracy of the envelope and instantaneous phase estimates obtained with the described techniques as a function of the group delay *D* we used the following metrics. All metrics are based on the ground truth envelope and instantaneous phase information extracted from the ground truth signal *s*[*n*] extracted non-causally from the real EEG data as described above and then shifted to match the specific delay value *D* of causal processing. We will denote the shifted ground truth envelope and phase sequences as *a*[*n − D*] and *ϕ*[*n − D*] correspondingly.

For performance assessment of envelope estimation methods, we calculated the Pearson correlation coefficient between estimated envelope *â*[*n*] obtained causally by each of the methods and the appropriately shifted ground truth envelope sequence *a*[*n − D*]:

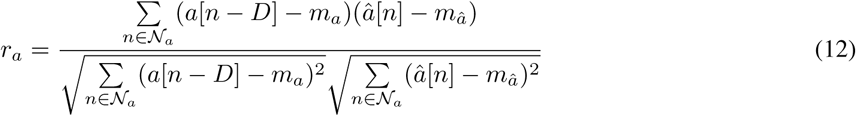

where 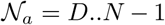 is the set of time indices with *N* = 30000, *m*_*a*_ and *m*_*â*_ are sample means averages of *a*[*n*] and *â*[*n*] over set *N*_*a*_

To asses performance of instantaneous phase estimation we used bias *b*_*ϕ*_, absolute bias |*b*_*ϕ*_| and the standard deviation *σ*_*ϕ*_ of the delayed ground truth phase *ϕ*[*n − D*] at the time moments when predicted 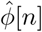 phase crosses 0. These metrics reflect the bias, absolute bias, and the variance in determining zero-crossing moments (negative-to-positive direction) of the delayed signal *s*[*n − D*].

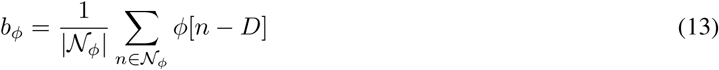

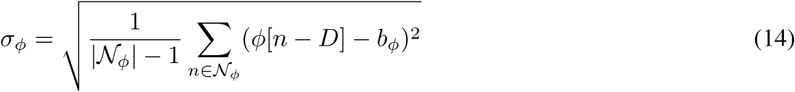

where 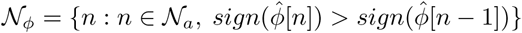 is the set of time moments when 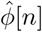 crosses 0.

#### Grid search procedure

To ensure that the compared methods operate optimally, for each of them, we defined grid search space as described in table 1 and, as described below, looked for the combination of parameters that ensured the best performance for each of the techniques. Here we use the following short method names: *rect* for envelope detector based on rectification of the band-filtered signal, *cfir* for the frequency domain least squares designed cFIR, *wcfir* for the frequency domain weighted least squares designed cFIR and *acfir* for RLS based cFIR filter update scheme. Note that sliding window narrow-band Hilbert transform approach (*hilb*) exactly matches the *cfir* on the range pf positive delays.

**Table 1:** Grid search space for each method.

| Method | Param. | Grid |
| --- | --- | --- |
| *rect* | $N_{bp}$ | 0, 5, ..., 100 |
| *cfir/hilb* | $N_t$ | 250, 500, 1000 |
| *wcfir* | $N_t$ | 250, 500, 1000 |
| *acfir* | $N_t$ | 250, 500, 1000 |
| | $N_u$ | 25, 50 |
| | $\mu$ | 0.7, 0.8, 0.9 |
| *ffiltar* | $p$ | 10, 25, 50 |
| | $N_e$ | 10, 25, 50 |

For each combination of parameters and fixed delay *D*, we computed the metrics defined above on the training set separately for each subject. Note that for *rect* and *hilb* no negative delay *D* is possible and for *ffiltar* we used only zero delay (*D* = 0) as this technique was formulated and used in the closed-loop experiments in this specific condition. Frequency band and weights for the *wcfir* approach were computed based on the same training data. We then used optimal values of parameters for each method corresponding to the maximum of *r*_*a*_, minimum for |*b*_*ϕ*_| and minimum of *σ*_*ϕ*_ values observed on the training set, and estimated the same performance metrics *r*_*a*_, *b*_*ϕ*_, |*b*_*ϕ*_| and *σ*_*ϕ*_ on the test data.

## 3 Results

Figure 3 shows the performance comparison results averaged over the data for ten subjects for the methods explored in this study. To ensure the best performance for each of the techniques, we used training data segments to tune each method’s parameters using the grid search procedure described in the Methods section. For each method in panels A, B, C and D, we show the performance metrics *r*_*a*_, *σ*_*ϕ*_, *b*_*ϕ*_, |*b*_*ϕ*_| reflecting envelope correlation accuracy (panel A), phase estimate variance (panel B), phase estimate bias (panel C) and phase estimate absolute bias (panel D) and described by equations 12,14 and 13 as observed on the test segments of the data with the optimal set of parameters identified over the independent training segments. For each such metric, the curves display their mean values averaged over 10 subjects as a function of the incurred delay. The error bars indicate the 95% confidence intervals obtained with 1,000 bootstrap iterations.

**Figure 3:**
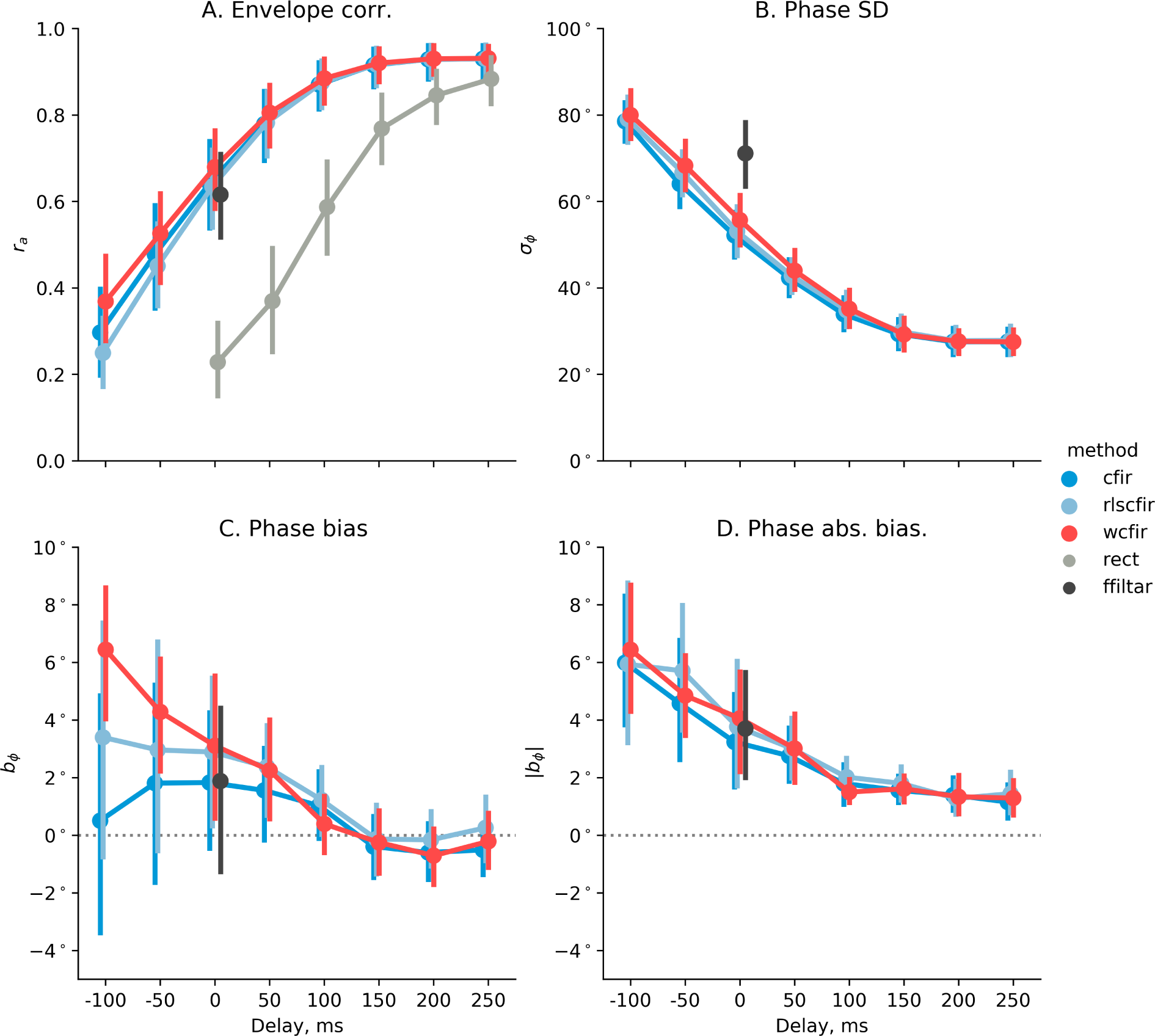
Four performance metrics vs incurred processing delay. A - dependence of the correlation coefficient *r*_*a*_ on delay *D* for the *cfir* family, for rectification based technique and AR-based extrapolation approach(*ffiltar*). B - phase estimation bias *b*_*ϕ*_ as a function of processing delay *D*. C - phase estimation variance *σ*_*ϕ*_ as a function of processing delay. D - phase estimation absolute bias |*b*_*ϕ*_| vs. *D*

As expected and according to Figure 3.A, the envelope estimation accuracy quantified by the correlation coefficient *r*_*a*_ deteriorates as the processing lag *D* decreases. The smaller the processing lag, the weaker is the correlation between the non-causally obtained ground-truth envelope and the envelope estimated in real-time by each of the methods. Noticeably, for all positive delays *rect* approach (gray line) has the worst performance that makes it practically unusable for latency values below 150 ms. Methods *cfir* (blue line), *wcfir* (red line) and *acfir* (light blue line) exhibit comparable accuracy. The approach described in [36] (*ffiltar*, black dot) that entails AR modelling of data and forward-backward filtering of the extended data chunk yields at zero latency the accuracy comparable to that delivered by the proposed *cfir* family of methods but requires specification of parameters describing AR model order, filter and extension window length and is significantly more intense computationally.

Panels B, C and D of Figure 3 show the results for real-time estimation of instantaneous phase as a function of time-lag *D*. For the positive delay values, the bias *b*_*ϕ*_ remains practically negligible and does not exceed 5^*◦*^ for all considered methods. Noteworthy, the *ffiltar* approach has been formulated for zero-latency and its phase estimation bias corresponding to *D* = 0 hovers around zero. However, as panel D demonstrates, the absolute bias of the phase estimate obtained by this technique is comparable to that delivered by the *cfir* family of methods and therefore close to zero values of the bias reflects symmetric around zero distribution of the bias observed in the 10 datasets explored. The methods from the *cfir* family exhibit a significant growth of phase estimation standard deviation (SD) as lag *D* decreases. Yet, at zero lag, the variance of the estimate delivered by the proposed here family of techniques appears to be significantly below that of the state of the art method introduced earlier in [36] and successfully used in the number of closed-loop stimulation studies. Here we reported the averaged observations made on the basis of the data recorded from 10 subjects as described in Section 2.3. The comparative performance of the explored methods may line-up differently for each of the datasets depending on the individual alpha-range SNR and other unaccounted factors.

This issue is clarified in Figure 4 that illustrates for individual subjects the same three performance measures as shown in Figure 3 but only for *D* = 0. Separate consideration for each subject allowed us to explore the performance as a function of SNR, which varied across individuals.

**Figure 4:**
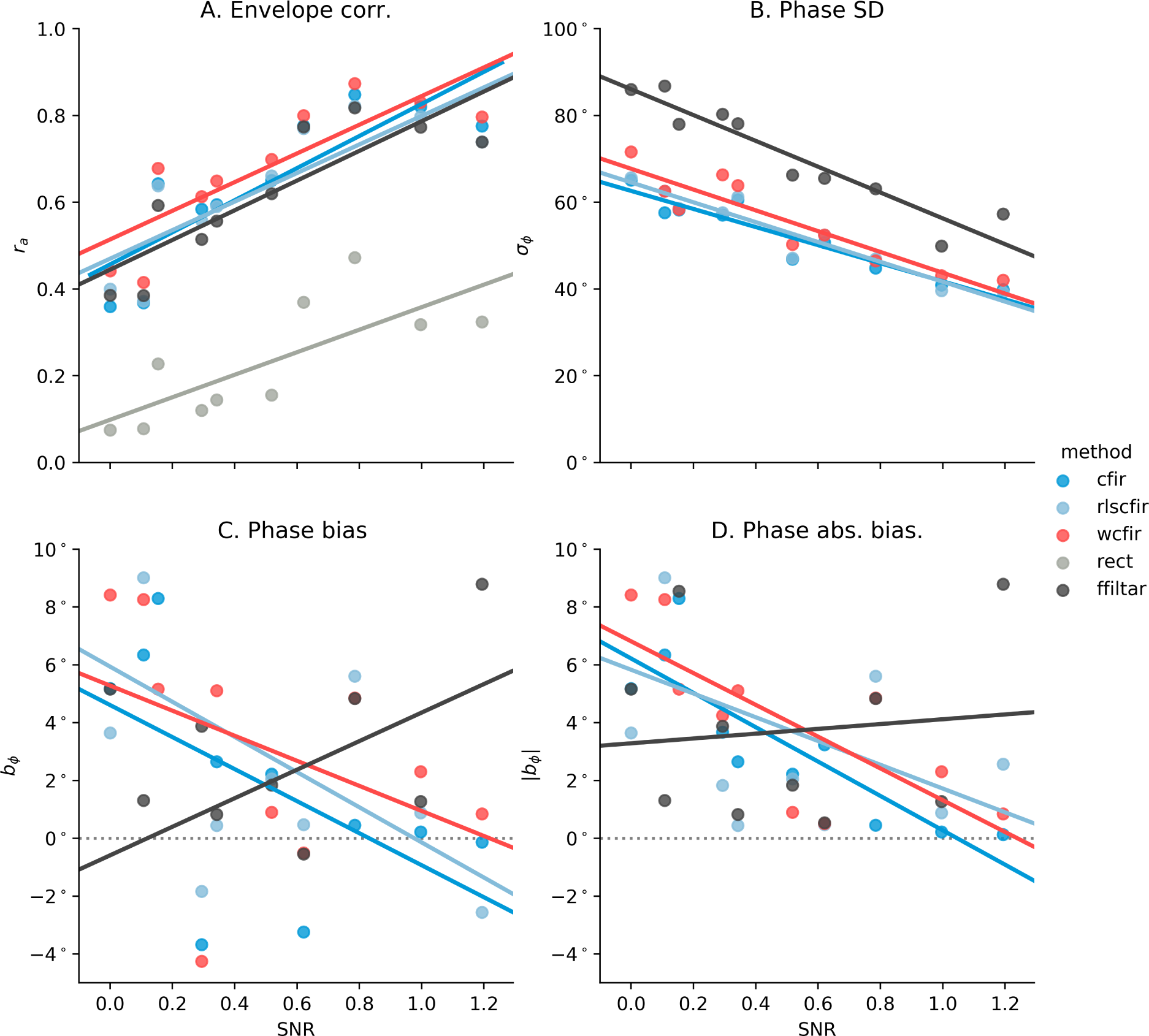
Four performance metrics vs. alpha rhythm SNR. A - envelope correlation coefficient *r*_*a*_, B - phase estimation standard deviation (SD) *σ*_*ϕ*_ and C,D - phase estimation bias *b*_*ϕ*_ and absolute bias |*b*_*ϕ*_|. Each dot corresponds to a dataset and is positioned on the x-axis according to the P4 alpha-rhythm SNR.

Figure 4 shows how the performance of the explored methods for *D* = 0 as a function of the SNR observed in each of the individual subjects. Metrics of the performance are the same as in Figure 3. For all methods envelope correlation *r*_*a*_ grows with SNR, panel A. Phase estimate standard deviation improves with SNR and appears to be consistently lower for the *cfir* family of methods. Phase estimate bias *b_ϕ_* shows positive skew for the low SNR value. The absolute bias for *ffiltar* technique appears to be independent of the SNR. For high SNR values starting with 1.0 the absolute bias value obtained by all methods from the *cfir* family appears to be consistently lower than that of the *ffiltar*.

In some versions of the closed-loop paradigms e.g. [24], a full-blown envelope reconstruction is not required. Instead, of interest is the discrete detection of time moments with high instantaneous band power. The transition into the zone of high instantaneous band power values may serve as a feedback or a trigger for either stimulus presentation or direct brain stimulation act. Suppose we want to perform detection of the time when the instantaneous rhythm power exceeds the 95% threshold. As shown in the left panel of Figure 5.A for the binary classification case the moments when the envelope falls into the top area above the dashed line are labeled as *High* and the rest of the moments are labeled as *Low*. The graph on the right panel of Figure 5.A shows balanced accuracy score (class recall average) for such binary detection task as a function of allowed processing delay parameter *D*. The analysis is done for one subject with median SNR selected from the pull of 10 subjects. We can observe that the best performance in the binary classification task is achieved by the weighted *cfir* method with zero processing latency (*D* = 0). Similar results for ternary classification of the three-state problem are shown in Figure 5.B. Just like in the binary case, the moments when the envelope falls into the top area corresponding to the 5% of the largest envelope values are labeled as *High* and additionally, label *Low* is assigned to the time instances when the envelope takes on values from the lowest 5%. The rest of the time moments are labeled as *Medium*. Interestingly, the *cfir* family of methods delivers the best performance for zero processing delay *D* = 0. In the binary classification scenario, we can achieve about 75% of balanced accuracy. The rectification-based approach at best provides just above 60% accuracy and that peaks in 100-150 ms processing lag range. The results are qualitatively the same for the ternary classification case.

**Figure 5:**
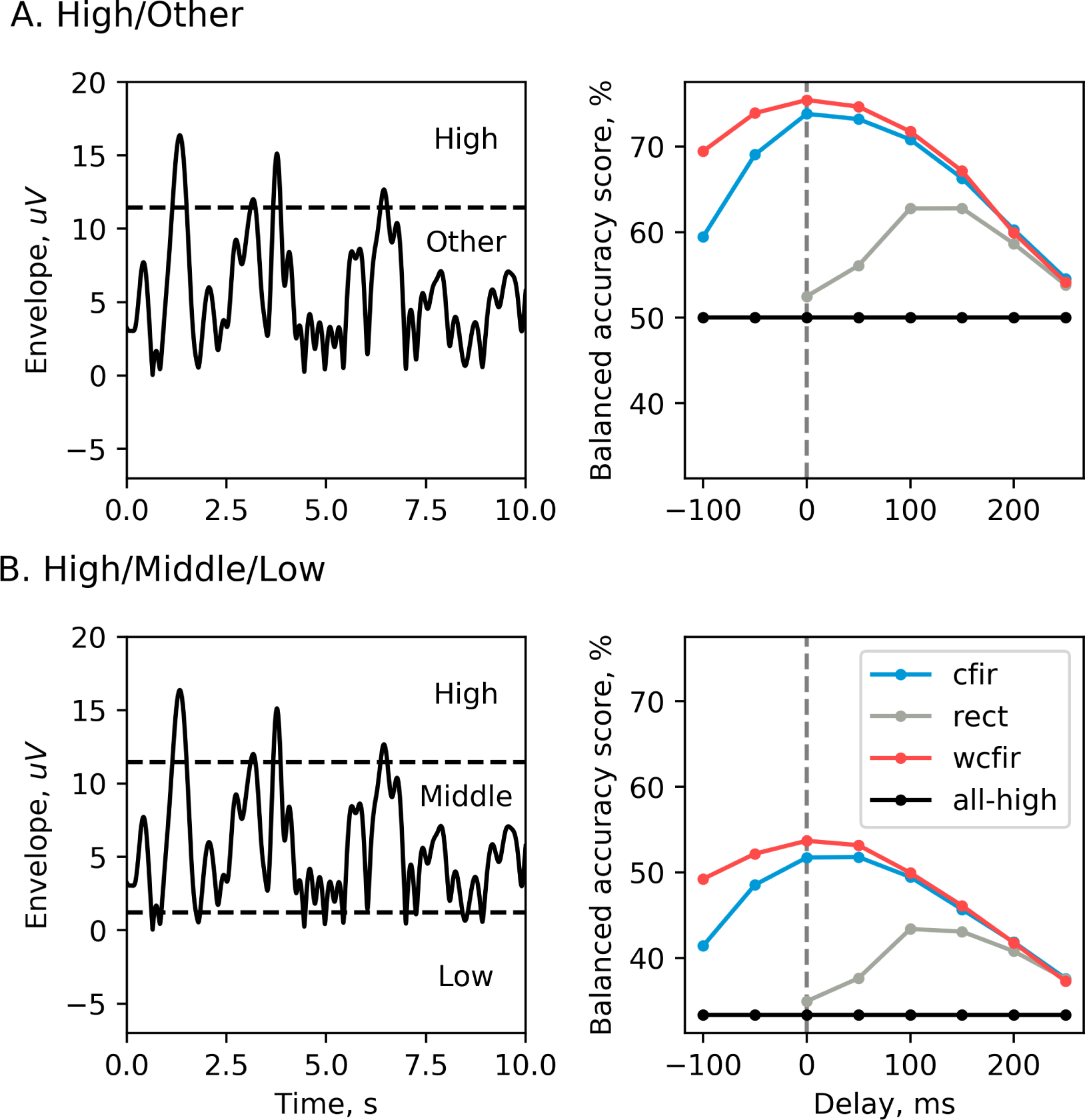
Discrete paradigm accuracy for one subject with median SNR. A) - binray classification task. The goal is to detect the time instances when alpha envelope is in the upper 5% quantile of its values. B) - ternary classification task to distinguish lower and upper 5% quantiles of the envelope values from the mid-range values falling into 5%-85% range.

Finally, to explore the morphology of the alpha-burst events in the *High*/*Other* classification task described above in Figure 6, we averaged the ground truth envelope around moments when the decoder crossed its own threshold. This computation was performed for *rect* and *wcfir* approaches for predefined delay parameters from [300, 100, 0, −100] ms set (for *rect* only positive values were used). Also, we computed averaged envelope across a set of randomly picked time moments (denoted as *rand*) and across moments when the ground truth envelope crossed the *High* threshold (denoted as *ideal*), which can not be done causally, see Figure 6.d

**Figure 6:**
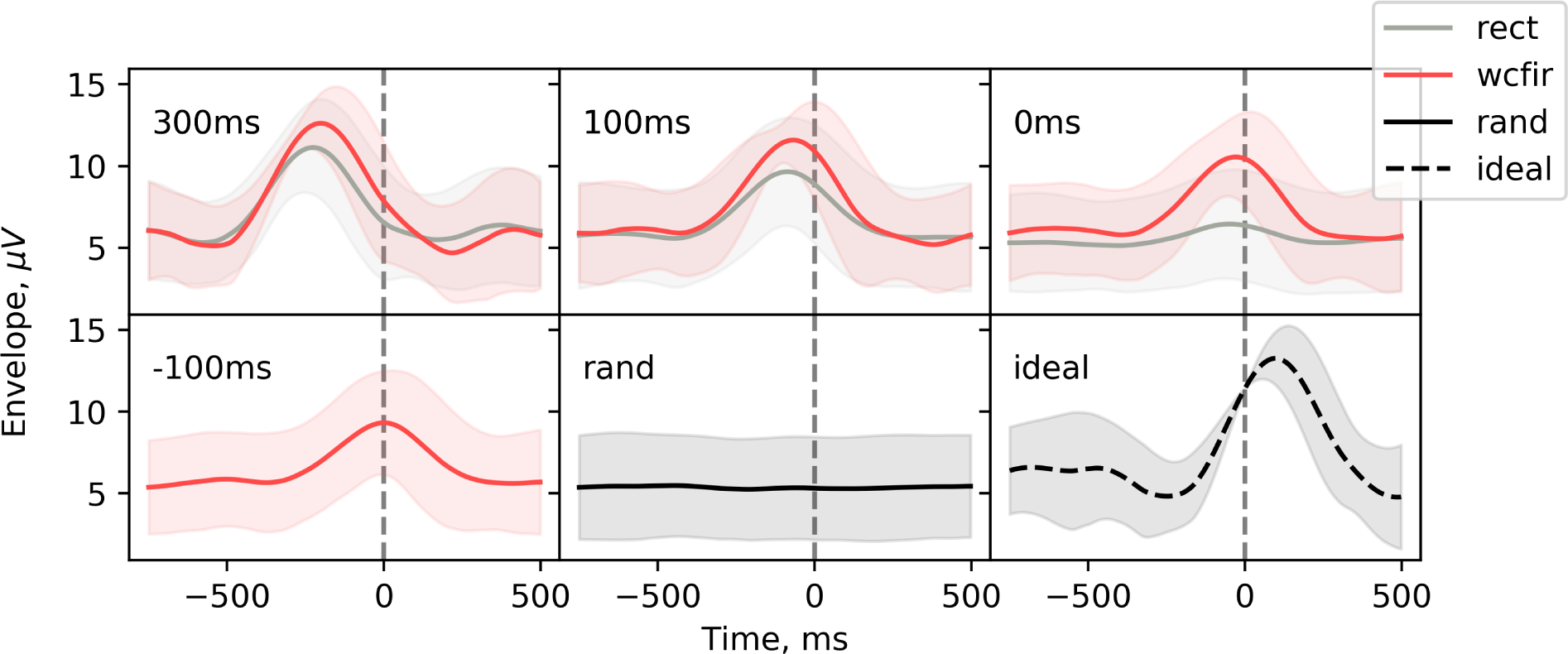
Average envelope computed using real-time detected markers of *High* events for different pre-specified processing delay values.

## 4 Discussion

The standard techniques for estimating instantaneous power of EEG rhythms, such as the methods based on the rectification of narrow-band filtered signal and the STFT based algorithms, incur significant delays, hindering the performance of BRC systems. Such delays, combined with the lags of acquisition hardware and the time required for stimulus presentation, result in significant lags between the actual brain activity and the signal used to control the experimental flow in BRC paradigms and/or utilized as a feedback to the subject. In the closed-loop stimulation paradigms, this would mean that the timing of the stimulating pulse can not be accurately aligned to the desired feature of the oscillatory brain activity. In NFB setting or settings requiring an explicit feedback signal that reflects subject’s performance, these standard approaches close the loop more than 300 ms past the targeted neural event [42]. Such delays may be especially harmful when the targeted brain rhythm patterns can be described as discrete events of a limited duration [7], [43],[44], where the feedback can arrive after the event has completed. Such low temporal specificity of the feedback signal hinders learning, especially in the automatic learning scenarios [30].

Here we systematically explored a series of methods for minimizing latency in RBC systems. We distinguished a family of best-performing techniques that are based on the least-squares filter design. These methods allow for a simpler and more transparent control over the accuracy–versus–latency trade-off compared to the other existing approaches.

Our results confirm that the proposed methodology based on least-squares filter design noticeably reduces latency of EEG envelope and phase estimation. This procedure is simple; yet its performance pars or exceeds that of the more complex approaches, such as the one based on the use of the AR-model [36], the most ubiquitous method for closed-loop studies [45], [18],[46],[47]. With our method, users can specify the desired delay and achieve the best possible envelope estimation accuracy possible with a linear method. As evident from Figure 3, the spectral density weighted cFIR technique allowed us to generate zero-latency feedback that accurately tracked the instantaneous power profile of the EEG-rhythm. The performance of our method on a typical segment of data can be appreciated from Figure 7 that shows true values and estimates of the envelope and phase obtained with cFIR method for various user specified lags.

**Figure 7:**
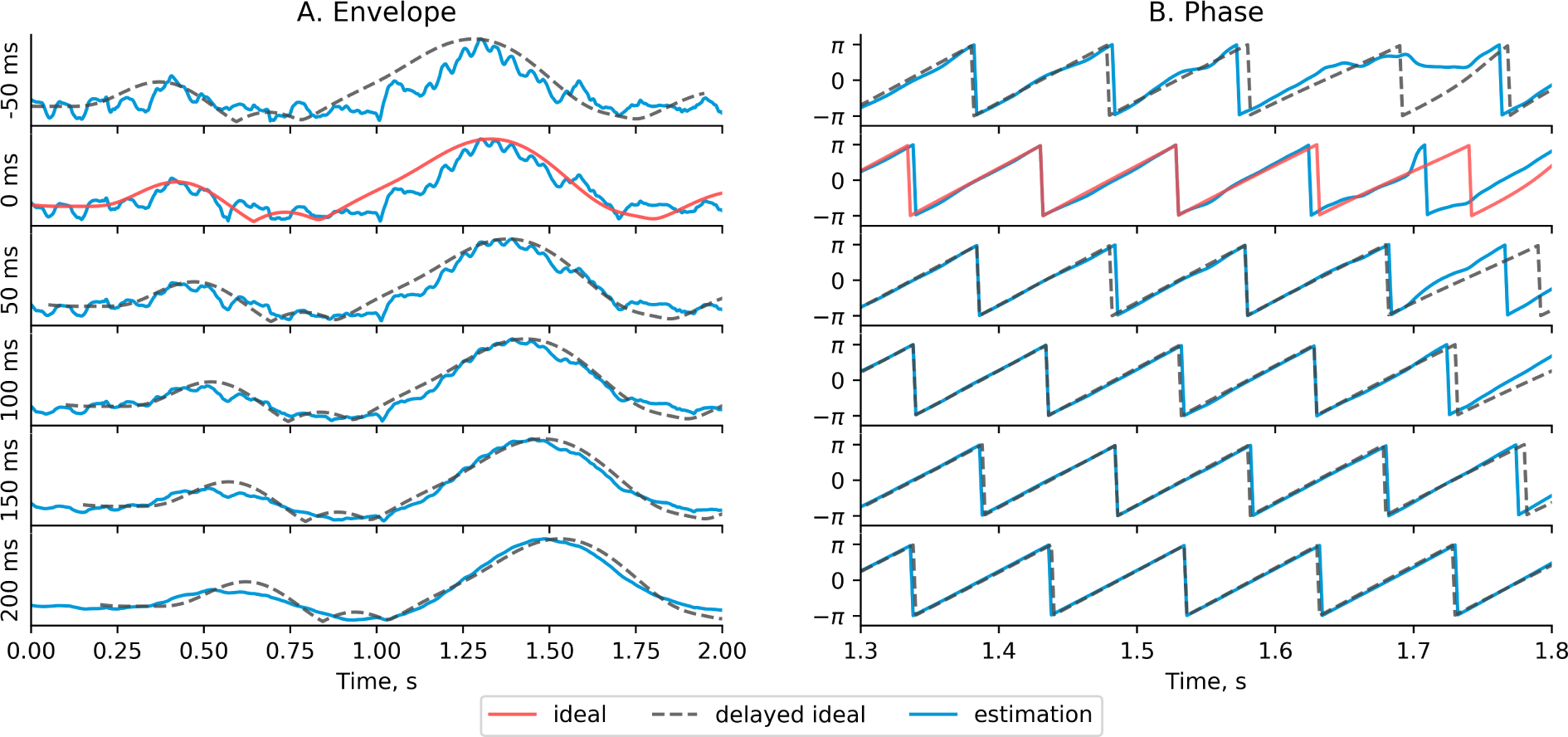
Envelope and phase estimates obtained by cFIR method (blue line) for different delay values *D*, each row corresponds to delays from −50ms to 200ms. Red line denotes ground-truth signal calculated by the ideal non-causal filter, gray dashed line denotes the same ground-truth signal but shifted by the corresponding delay *D* to facilitate comparison

We see this work as a systematic effort aimed at building a zero- or even negative-latency feedback systems that will allow transferring the predictive control methodology successfully exercised in technical systems to the tasks where the brain is the controlled object [48]. As shown in 5, *wcFIR* approach allows for correct forward prediction 100 ms ahead of rhythmic activity bursts, with AUC exceeding 70%. This illustration of the successful predictive behavior suggests that the proposed family of simple approaches together with the necessary hardware optimization will open up a way for the implementation of predictive control that enables a more efficient interaction with the functioning brain.

While our method reduces feedback latency, this reduction comes at a cost of less accurate envelope estimation. Deterioration of performance is especially sizeable in when the SNR is low and therefore, for the latency-reduction algorithm to be efficient, care should be taken to improve the SNR with such methods as spatial filtering of multi-electrode recordings.

The optimal latency-accuracy trade-off is the issue that needs to be addressed for each particular application and each particular subject. As shown in Figure 3, the methods outlined in this work allow the users to smoothly control this trade-off and choose the optimal operational point for each specific application.

As mentioned above, to achieve the true predictive scenario, though, the improvements need to be made not only of the signal processing algorithms but also of the hardware employed for signal acquisition, as well as the low-level software that handles EEG data transfer from the acquisition device to the computer memory buffer. To this end, it is worth considering specialized systems based on the FPGA programmable devices that eliminate the uncertain processing delays present in computer operating systems not designed to operate in real-time.

In the context of neurofeedback, additional consideration should be given to the physiological aspects of the sensory modality used to deliver the feedback signal. For instance, it is known that visual inputs, although very informative [33], are processed slower compared to tactile inputs and therefore tactile feedback could be a better option for predictive control.

The signal processing approaches presented here could be advanced by employing more sophisticated decision rules capable of extracting the hidden structure from the data. Thus, convolutional neural networks [49],[50] and novel recursive architectures hold a significant promise to further improve the accuracy of real-time zero-lag envelope and phase estimation.

## Acknowledgments

This work is supported by the Center for Bioelectric Interfaces NRU HSE, RF Government grant, ag. No.14.641.31.0003. We also thank Dr. Maria Volodina for the careful proofread of the final version of this manuscript.

